# Biochemical analyses reveal amino acid residues critical for cell cycle-dependent phosphorylation of human Cdc14A phosphatase by cyclin-dependent kinase 1

**DOI:** 10.1101/242016

**Authors:** Sara Ovejero, Patricia Ayala, Marcos Malumbres, Felipe X. Pimentel-Muiños, Avelino Bueno, María P. Sacristán

**Affiliations:** Instituto de Biología Molecular y Celular del Cáncer (IBMCC), Universidad de Salamanca-CSIC, Campus Miguel de Unamuno, 37007 Salamanca, Spain.; Departamento de Microbiología y Genética, Universidad de Salamanca, Campus Miguel de Unamuno, 37007 Salamanca, Spain.; Current affiliation: Institute of Human Genetics, CNRS UMR 9002, CNRS, Université de Montpellier, 34396 Montpellier, France.; Centro Nacional de Investigaciones Oncológicas (CNIO), E-28029 Madrid, Spain.

## Abstract

Cdc14 enzymes compose a family of highly conserved phosphatases that are present in a wide range of organisms, including yeast and humans, and that preferentially reverse the phosphorylation of Cyclin-Dependent Kinase (Cdk) substrates. The budding yeast Cdc14 orthologue has essential functions in the control of late mitosis and cytokinesis. In mammals, however, the two Cdc14 homologues, Cdc14A and Cdc14B, do not play a prominent role in controlling late mitotic events, suggesting that some Cdc14 functions are not conserved across species. Moreover, in yeast, Cdc14 is regulated by changes in its subcellular location and by phosphorylation events. In contrast, little is known about the regulation of human Cdc14 phosphatases. Here, we have studied how the human Cdc14A orthologue is regulated during the cell cycle. We found that Cdc14A is phosphorylated on Ser411, Ser453 and Ser549 by Cdk1 early in mitosis and becomes dephosphorylated during late mitotic stages. Interestingly, *in vivo* and *in vitro* experiments revealed that, unlike in yeast, Cdk1-mediated phosphorylation of human Cdc14A did not control its catalytic activity but likely modulated its interaction with other proteins in early mitosis. These findings point to differences in Cdk1-mediated mechanisms of regulation between human and yeast Cdc14 orthologues.

## Introduction

Cdc14 family members are dual-specificity phosphatases that preferentially reverse Cdk-dependent phosphorylations ^1^. They are highly conserved and are present in eukaryotes ranging from yeast to mammals. Their functions are quite well established in yeast. In *Saccharomyces cerevisiae*, Cdc14 exerts an essential role in regulating late mitotic events, mitotic exit and cytokinesis by reversing phosphorylation of different CDK substrates ^2–5^. In the fission yeast *Schizosaccharomyces pombe*, the Cdc14 orthologue, Flp1/Clp1 (referred to as Flp1 hereafter), although a non-essential gene, also targets mitotic cyclin-dependent Cdk1 substrates to regulate late mitotic events and cytokinesis ^6–10^. Moreover, both budding and fission yeast Cdc14 orthologues have been involved in DNA damage response mechanisms ^11–13^. Human Cdc14 phosphatases, Cdc14A and Cdc14B, although able to rescue Cdc14-deficient yeast cells suggesting conserved functions ^14,15^, do not play a prominent role in controlling late mitotic events ^16^. Mammalian Cdc14s have been involved in several cellular processes such as the centrosome duplication cycle, control of Cdk1 activity, cytokinesis, cell adhesion and migration, transcription regulation and also DNA damage response ^16–29^.

All Cdc14 phosphatases share a highly conserved N-terminal domain, which comprises the PTP (protein-tyrosine phosphatase) catalytic motif, and have a variable C-terminal domain, whose length and sequence conservation differs among the different species. The carboxyl terminus contains a nuclear export sequence and has been shown to be subjected to post-translational modification ^11,30^. Regulation of Cdc14 is also fairly well characterized in yeast. In *S. cerevisiae* the activity of Cdc14 is largely controlled at the level of subcellular localization. Thus, Cdc14 is maintained in a nucleolar-bound inactive form during interphase and in a nucleolar-released active state during late mitosis. Cdc14 nucleolar release and activation starts at the onset of anaphase, the time at which Cdc14 initiates essential roles for nuclear and cytoplasmic divisions, and are promoted by the coordinated and consecutive action of the mitotic networks FEAR (fourteen early anaphase release) and MEN (mitotic exit network) ^4,31,32^. In the fission yeast *S. pombe*, Flp1 regulation also involves a change in its subcellular localization. Thus, the interphase nucleolar Flp1 is released from the nucleolus early in mitosis to concentrate on the kinetochores and contractile ring and to disperse throughout the nucleus and cytoplasm ^9,10,33^. But in this case, an additional mechanism controls its catalytic activity. Flp1 is phosphorylated by Cdk1 during early mitosis to stay inactive until mitotic exit, the time at which the protein is activated by autodephosphorylation to participate in the orderly dephosphorylation of Cdk1 substrates ^30^.

As in yeast, human Cdc14 phosphatases have different localizations throughout the cell cycle. Thus, Cdc14A and Cdc14B, concentrated in the centrosomes and nucleolus, respectively, during interphase, become dispersed throughout the cell upon entry into mitosis ^18,29^. We have previously shown that Cdc14A modulates the timing of mitotic entry through the regulation of both positive and negative Cdk1 regulators, Cdc25B phosphatase and Wee1 kinase, respectively ^26,28^. Cdc14A has also been involved in late mitotic processes, such as chromosome segregation, and later on, cytokinesis ^18,34,35^. These observations suggest that Cdc14A phosphatase participates in the dynamic control of protein phosphorylation during mitosis, and that it should therefore be subjected to strict spatiotemporal regulation.

Here, we describe mitotic-specific phosphorylation of human Cdc14A by Cdk1-Cyclin B1 complexes. Cdc14A gets hyperphosphorylated during early mitosis and then, at the same time as Cdk1 inactivation during late mitosis, Cdc14A becomes dephosphorylated. In addition, we discovered that although Cdc14A has autodephosphorylation capacity, its dephosphorylation during mitotic exit is regulated by other phosphatases. Moreover, we found that Cdk1-mediated Cdc14A phosphorylation does not regulate either its catalytic activity (in contrast to what has been observed in yeast) or its subcellular localization or stability. However, Cdk1-mediated Cdc14A phosphorylation in early mitosis may modulate its protein interaction pattern. These results suggest a clear divergence between yeast and human Cdc14 phosphatases, regarding to the mechanisms of their regulation through the cell cycle.

## Results

### Human Cdc14A is a phosphoprotein with autodephosphorylation activity

Based on the banding pattern obtained by immunodetection, it has been suggested that human Cdc14A phosphatase could be a phosphoprotein ^29^. When ectopically expressed, we routinely noticed that electrophoretic mobility of the inactive form of Cdc14A, phosphatase dead or Cdc14A(PD), appeared slightly decreased when compared with the wild-type protein (Supplementary Figure S1), suggesting that Cdc14A is in fact phosphorylated in the cell and that it is able to modify its own phosphorylation state. To confirm this observation, HEK293T cells ectopically expressing Flag-Cdc14A wt or Flag-Cdc14A(PD) were treated with okadaic acid (OA), an inhibitor of two of the main broad-specificity protein phosphatases, PP1 and PP2A, to enhance the phosphorylation of multiple cellular proteins, and Cdc14A was analysed by immunoblotting using phosphate-affinity SDS-PAGE gels (Phos-tag gels). As shown in Fig. 1a, the treatment with OA produced a strong up-shift of the active form of Cdc14A, confirming that it is a phosphoprotein. In addition, we observed that the migration of the inactive form was in fact slower than that of the active form in both untreated and OA treated cells, demonstrating the ability of Cdc14A to act on itself, even under high kinase activity. These results also demonstrate that Cdc14A is an okadaic acid insensitive phosphatase. The treatment of cell extracts with lambda protein phosphatase restored the mobility of Flag-Cdc14A wt and Flag-Cdc14A(PD) to the faster migrating band, also demonstrating that the observed change in the electrophoretic mobility of Cdc14A corresponded to the phosphorylated forms of the protein (Fig. 1b).

**Figure 1.**
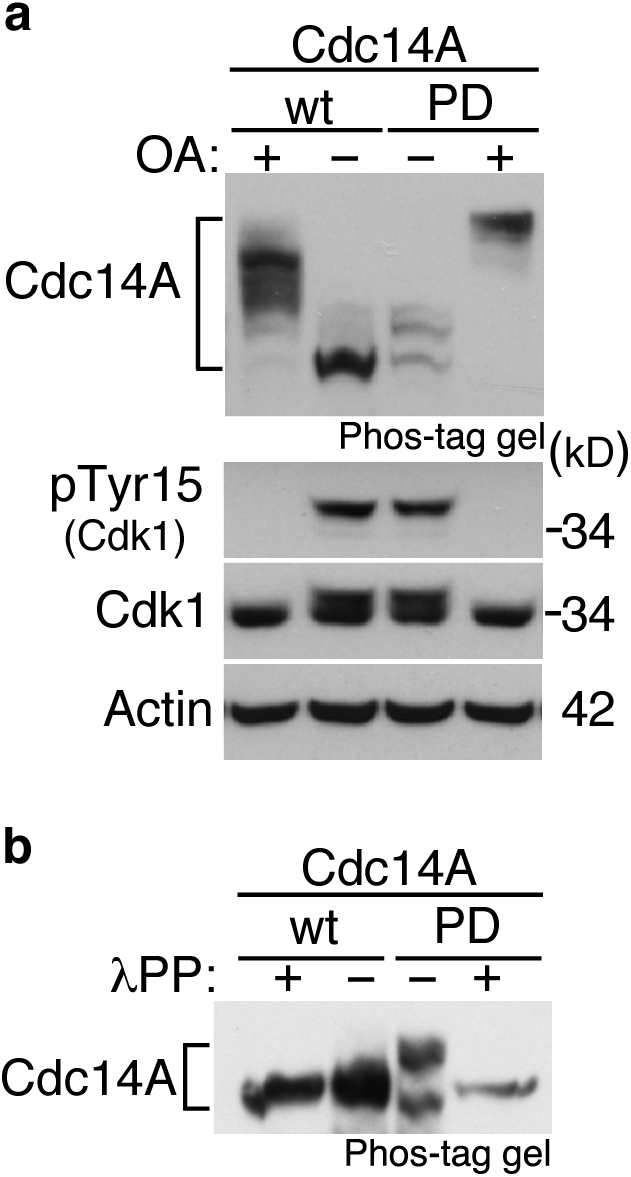
Cdc14A is a phosphoprotein with autodephosphorylation ability. **(a)** HEK293T were transiently transfected to express Flag-Cdc14A or its inactive form Flag-Cdc14A(PD). After 24 hours, cells were treated with OA (0,5 µM), or not, during 2 hours and cellular extracts were obtained and analyzed by immunoblot against the indicated proteins. Activation of Cdk1, as a consequence of OA treatment, was confirmed by phospho-Cdk1(Tyr15) detection. Phos-tag gel was used to specifically identify the phosphorylation status of the Cdc14A forms. **(b)** Total protein extracts from HEK293T cells transfected with Flag-Cdc14A or Flag-Cdc14A(PD), as described in (a), were incubated with or without lambda phosphatase (λPP), resolved by Phos-tag gels and analyzed by immunoblot with anti-Cdc14A antibodies.

### Cdk1 phosphorylates Cdc14A during early mitosis

To gain insight into the phosphorylation of Cdc14A, we studied its phosphorylation state during the cell cycle by immunoblotting and Phos-tag gel analysis. Since available Cdc14A antibodies did not detect the endogenous protein, we generated U-2-OS-HA-Cdc14A and U2OS–HA-Cdc14A(PD) stable cell lines expressing low levels of the proteins. Ectopic expression of these proteins did not have any effect on cell cycle progression, or on its specific subcellular localization (Supplementary Figure S1). U-2-OS-HA-Cdc14A cells were synchronized at the G1/S transition or at early mitosis by a double thymidine block or nocodazole treatment, respectively, and then released into fresh medium to allow progression through the cell cycle. We observed that Cdc14A becomes hyperphosphorylated at entry into mitosis, maintains this state of phosphorylation during early mitosis and is then dephosphorylated as cells exit from mitosis (Fig. 2a). Phosphorylation of Cdc14A was also observed in mitotic cells not treated with nocodazole (Supplementary Figure S1), which excludes the possibility that Cdc14A phosphorylation could be a consequence of the nocodazole induced-SAC (Spindle Assembly Checkpoint) activation. These results indicate that Cdc14A is phosphorylated during early mitosis and dephosphorylated at mitotic exit.

**Figure 2.**
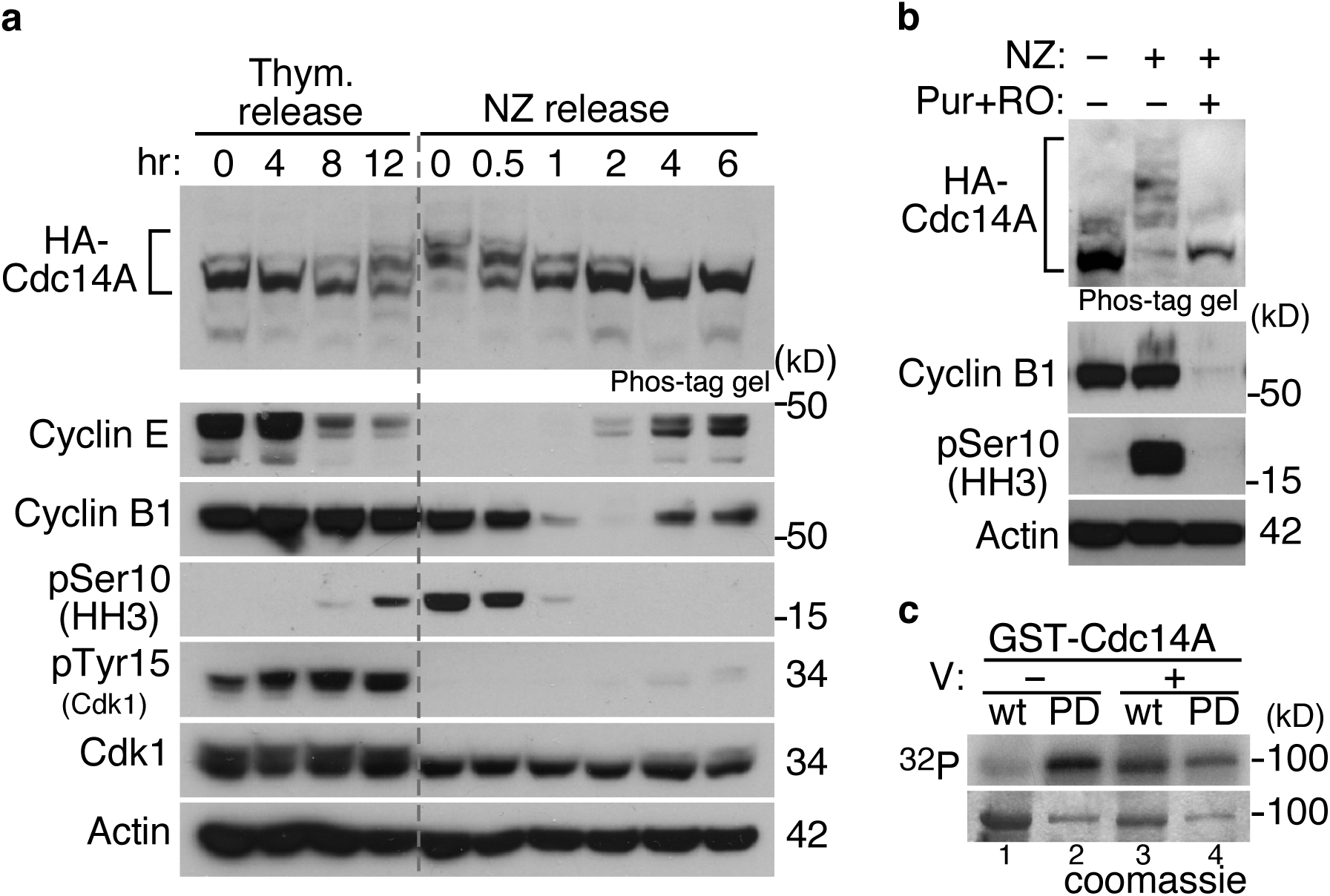
Cdk1-dependent phosphorylation of Cdc14A early in mitosis. **(a)** U-2-OS cells stably expressing retrovirally transduced HA-Cdc14A were synchronized at G1/S by a double thymidine treatment (0 hr) and then released into fresh medium containing nocodazole (NZ) to allow progression through G2 phase and G2/M transition and to avoid mitotic exit. For mitosis-synchronized cells, the cultures were treated with nocodazole during 12 hours and rounded mitotic cells were collected (0 hr) or released into fresh medium to allow progression through mitosis and G1 phase. Phosphorylation state of Cdc14A was analyzed by immunoblot, using Phos-tag gels, at the indicated time points. Progression through the cell cycle was monitored by immunoblot against the indicated proteins. **(b)** U-2-OS-HA-Cdc14A cells were treated with nocodazole during 12 hours. Rounded mitotic cells were selected by shake-off and then treated with the vehicle dimethyl sulfoxide or the Cdk1 inhibitors Purvalanol A (Pur) and RO-3306 (RO) during 3,5 hours. Samples were analyzed by immunoblot with the indicated antibodies. Asynchronous cells were also analyzed, and phosphorylation state of HA-Cdc14A was checked using Phos-tag gels. **(c)** GST-Cdc14A, (wt), and GST-Cdc14A(PD), (PD), were phosphorylated *in vitro* by Cdk1-Cyclin B1 complexes in the presence (+) or absence (−) of 1mM sodium orthovanadate (V), using γ(^32^P)ATP. Reactions were resolved by SDS-PAGE and analyzed by coomassie staining and autoradiography. These results show representative data from al least three different experiments.

It has been shown that the fission yeast Cdc14 orthologue, Flp1, is phosphorylated by Cdk1 during early mitosis^8,9,30^. To check whether Cdk1 is the mitotic kinase responsible for Cdc14A phosphorylation, nocodazole arrested cells were treated with the Cdk1 inhibitors Purvalanol and RO3306, and Cdc14A phosphorylation state was analyzed by Phos-tag gel. As shown in Fig. 2b, inhibition of Cdk1 fully abolished the mitotic band shift, suggesting that hyperphosphorylation of Cdc14A is Cdk1-dependent. To confirm that Cdk1 phosphorylates Cdc14A, both GST-Cdc14A or GST-Cdc14A(PD) recombinant proteins, purified from *E. coli*, were subjected to *in vitro* kinase assays using Cdk1-Cyclin B1 complexes. GST-Cdc14A proteins were detected as radiolabelled proteins (Fig. 2c), demonstrating that Cdc14A is a substrate of Cdk1. Interestingly, incubation of Cdc14A(PD) with Cdk1 caused increased radiolabel incorporation compared with wild-type Cdc14A acting as substrate (Fig. 2c, lanes 1 and 2). This suggests that Cdc14A autodephosphorylates during the reaction. The addition of an inhibitor of tyrosine family phosphatases, sodium orthovanadate, to the kinase reactions yielded similar radiolabel incorporations in both the wild-type and the inactive Cdc14A forms (Fig. 2c, lanes 3 and 4), indicating that Cdc14A has autocatalytic activity. Together, these data demonstrate that Cdk1 phosphorylates Cdc14A during early mitosis, and that Cdc14A has the capacity to reverse, at least partially, its own Cdk1-mediated phosphorylation *in vitro*.

### Cdc14A dephosphorylation at the exit from mitosis is not only due to autoregulation

Since Cdk1 phosphorylates Cdc14A during early mitosis and Cdc14A has autophosphatase activity, we surmised that Cdc14A controlled its own dephosphorylation at the exit from mitosis as Flpl does in the fission yeast *S. pombe* ^30^. In order to address this issue, we examined the phosphorylation state of its inactive form throughout the cell cycle, using the U2OS–HA-Cdc14A(PD) cell line. As expected, Cdc14A(PD) showed higher phosphorylation levels than the active form at all time points analysed (Fig. 3a, and compare with Fig. 2a). Moreover, the inactive form gets hyperphosphorylated at entry into mitosis and becomes dephosphorylated with similar kinetics to that of the active phosphatase, suggesting that Cdc14A does not control its full dephosphorylation during mitotic exit (Fig. 3a). Since U-2-OS-HA-Cdc14A(PD) cells also express endogenous Cdc14A, we could not exclude the possibility that the observed dephosphorylation was carried out by the endogenous protein. In fact, *in vitro* phosphatase assays revealed that Cdc14A, and not the Cdc14B homologue, was able to remove Cdk1-mediated phosphorylations from Cdc14A(PD), showing its ability to autodephosphorylate *in trans* (Fig. 3b,c). In order to test whether or not endogenous Cdc14A phosphatase activity was responsible for Cdc14A(PD) dephosphorylation in these cells, we generated RPE1 and RPE1-Cdc14A^-/-^ ^16^ cell lines stably expressing retrovirally transduced HA-Cdc14A(PD) at low but detectable levels. Cells were synchronized in mitosis by nocodazole treatment and mitotic cells were collected by shake off and released into fresh medium to allow progression through mitosis. As shown in the Fig. 3d, in both wild-type RPE1 or Cdc14A deficient RPE1 cells, ectopic Cdc14A(PD) protein appeared dephosphorylated at the exit from mitosis, indicating that endogenous Cdc14A does not revert its own hyperphosphorylation at mitotic exit. PP2A phosphatase has been shown to be involved in the dephosphorylation of Cdk1 substrates at mitotic exit ^36–38^. We found that Cdc14A interacts with PP2A α-catalytic subunit (Supplementary Figure S2), suggesting that PP2A could be the phosphatase responsible for mitotic Cdc14A dephosphorylation. These results show that although human Cdc14A is a Cdk1 substrate at early mitosis and has autodephosphorylation capacity, it does not share with its orthologue Flp1 the full capacity to remove its own Cdk1-mediated phosphorylation at late mitosis.

**Figure 3.**
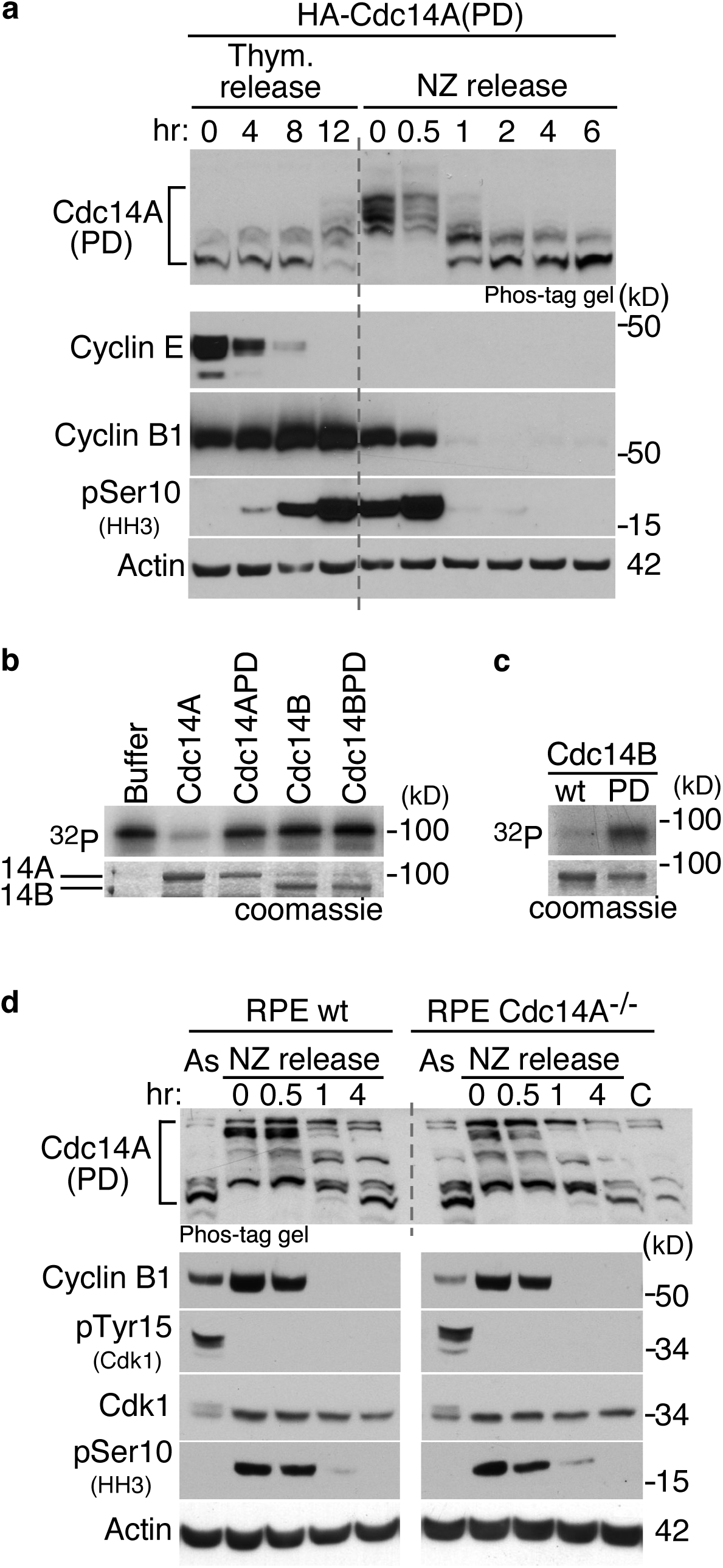
Cdc14A does not autodephosphorylate at the exit from mitosis. **(a)** U-2-OS cells stably expressing the retrovirally transduced inactive form HA-Cdc14A(PD) were synchronized as indicated in Fig. 2a. Phosphorylation state of the inactive form of Cdc14A was analyzed by immunoblot, using Phos-tag gels, at the indicated time points. Progression through the cell cycle was monitored by immunoblot against the indicated proteins. **(b)** GST-Cdc14A(PD) was phosphorylated *in vitro* by Cdk1-Cyclin B1 complexes in the presence of γ(^32^P)ATP. Samples were then washed and incubated with buffer alone or with 100ng of GST-Cdc14A, GST-Cdc14A(PD), GST-Cdc14B or GST-Cdc14B(PD). **(c)** GST-Cdc14B, (wt), and GST-Cdc14B(PD), (PD), were phosphorylated *in vitro* by Cdk1-Cyclin B1 complexes in the presence of γ(^32^P)ATP and without sodium orthovanadate. Note that incubation of Cdc14B(PD) with Cdk1 yielded increased radiolabel incorporation relative to wild-type Cdc14B acting as substrate, which demonstrates the activity of GST-Cdc14B. These results show representative data from at least three different experiments. **(d)** wild-type and Cdc14A knockout RPE cells (Cdc14A^-/-^) stably expressing retrovirally transduced HA-Cdc14A(PD) were treated with nocodazole during 12 hours. Rounded mitotic cells were collected (0 hr) and released into fresh medium to allow progression through mitosis. Phosphorylation state of Cdc14A(PD) was analyzed by Phos-tag gels at the indicated time points. Progression through the cell cycle was monitored by immunoblotting against the indicated proteins. C: RPE Cdc14A^-/-^ control extracts used to identify specific Cdc14A bands. As: Asynchronous cultures.

### Identification of Cdk1-specific phosphosites on Cdc14A

Our next aim was to identify the mitotic phosphorylation sites on Cdc14A. Sequence analysis showed that Cdc14A possesses seven minimal consensus motifs (S-T/P) for Cdk1 phosphorylation (Fig. 4a); two of them located in the N-terminal domain (ND) and five within the C-terminal domain (CD). We analysed two truncated versions of Cdc14A, the highly evolutionary conserved amino-terminal domain (Cdc14A-ND) and the carboxyl regulatory domain (Cdc14A-CD) (Fig. 4a), ectopically expressed in HEK293T cells. Nocodazole and OA treatments did not affect the electrophoretic mobility of Cdc14A-ND. On the contrary, Cdc14A-CD showed a marked delay under OA treatment conditions (Fig. 4b), suggesting that phosphorylation occurs on one or several residues of the C-terminal domain. However, since the absence of the band shift observed for Cdc14A-ND did not definitively exclude that phosphorylation events had not taken place on the two potential Cdk1 phosphorylation sites lying within that domain, we sough to analyse the full-length Cdc14A by mass spectrometry. Myc-tagged Cdc14A or Cdc14A(PD) were expressed in HEK293T cells and purified by immunoprecipitation from cells asynchronously growing or treated with OA to increase Cdk1 kinase activity. Cdc14A immunoprecipitates were resolved by SDS-PAGE and the bands corresponding to Cdc14A proteins were digested with proteases. The resultant peptides were analyzed by SIMAC (sequential elution from IMAC, ^39^) and MALDI-TOF. Three serine phosphorylation residues corresponding to minimum Cdk phosphorylation consensus sites and located at the carboxyl terminus were identified (S411, S453 and S549; Fig. 4a,c). Neither of the two potential Cdk1 residues lying on the amino terminal of Cdc14A appeared modified in the analysis, which was in agreement with our previous results (Fig. 4b). To further confirm the phosphorylation of these serines, we transfected HEK293T cells with the corresponding single, double or triple nonphosphorylatable serine to alanine full-length Cdc14A mutants and checked their phosphorylation state by Phos-tag gels after nocodazole treatment. Since we had previously shown that Cdc14A was able to partially autodephosphorylate even at high kinase activity conditions (Fig. 1a), which could interfere with the results, the analysis was performed using the inactive form of the protein. As shown in Fig. 4d, the mutation of single Ser411, Ser453 or Ser549 to nonphosphorylatable Ala residues partially modified the phosphorylation pattern of Cdc14A. Moreover, the triple mutation S411/453/549A fully abolished the band shift, causing the molecule to run with the same mobility as the wild-type protein from asynchronously growing cells, suggesting that Cdc14A is phosphorylated on those three residues.

**Figure 4.**
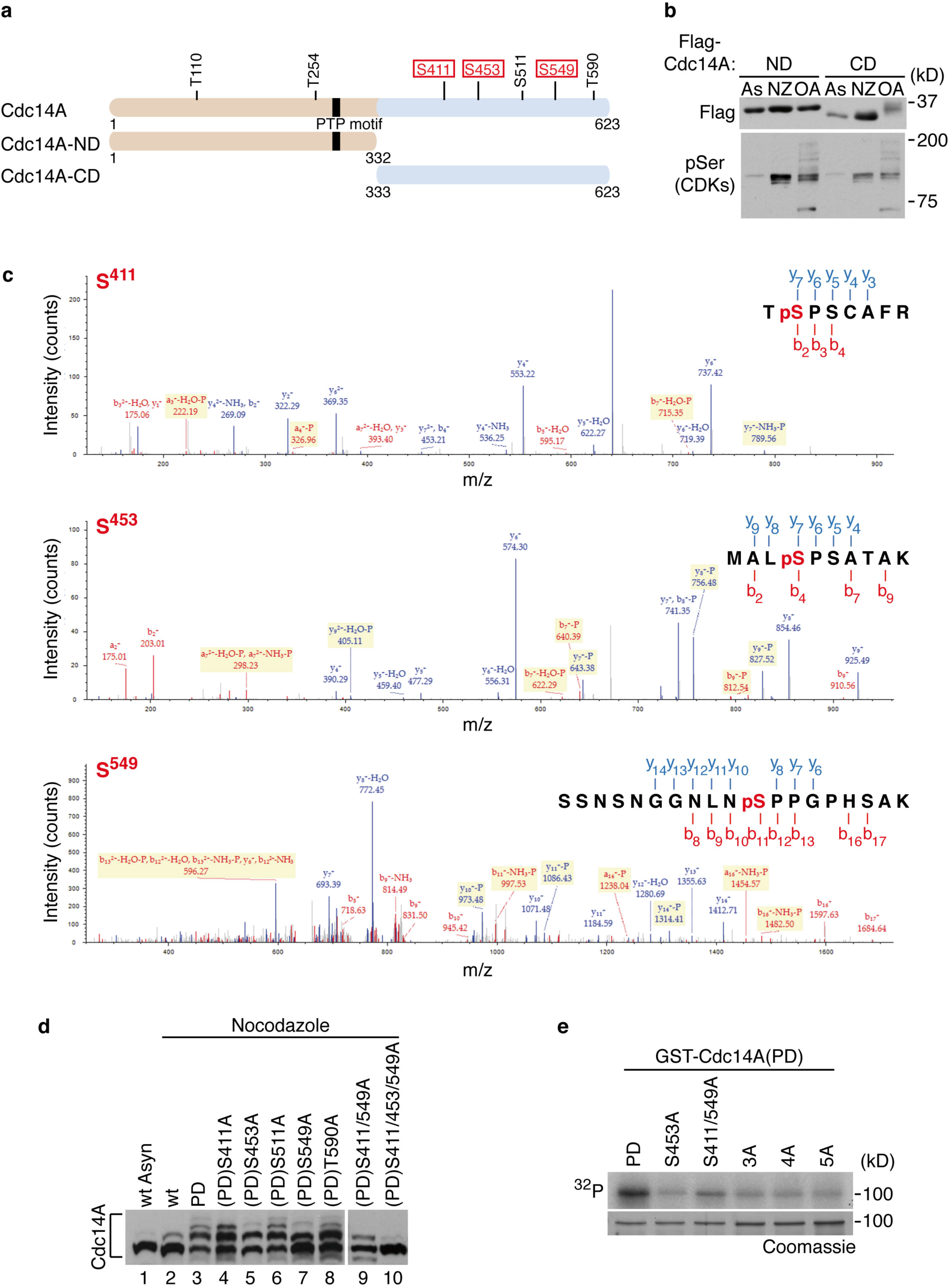
Cdc14A is phosphorylated by Cdk1 at Ser411, Ser453 and Ser549. **(a)** Schematic representation of full-length Cdc14A protein showing the distribution of all CDK phosphorylation consensus sites (S/T-P). Boxed sites represent those identified by mass spectrometric analysis of Cdc14A immunoprecipitates from HEK293T cells treated with OA. The amino-terminal and carboxy-terminal Cdc14A constructs are also indicated. **(b)** HEK293T cells were transfected with Flag-tagged N-terminal (ND; aa 1–332) or C-terminal (CD; aa 333–623) domains of Cdc14A. After 12 or 24 hours of transfection cells were treated with nocodazole (NZ, 50 ng/ml, 12 hours) or OA (0.5 µM, 2 hours), respectively. As: asynchronous, untreated cells. Cdc14A fragments were detected by immunoblotting with anti-Flag antibodies. The detection of phospho-Ser of Cdk substrates was used to confirm the efficiency of both NZ and OA treatments. Note that phosphorylation levels were higher in treated samples compared with non-treated asynchronous (As) cellular extracts. **(c)** Tandem mass spectrometry (MS/MS) spectra for Cdc14A phosphopeptides obtained by multistage activation CID. Representative MS/MS spectra show fragmentation of the peptides: TpSPSCAFR, MALpSPSATAK and SSNSNGGNLNpSPPGPHSAK. The peptide sequence above each representative spectrum shows theoretical “b” and “a” ion identifications (red, NH2-terminal fragments) and theoretical “y” ion identifications (blue, COOH-terminal fragments). Peaks in the spectrum that are marked red correspond to matched “b” and “a” ions, and peaks that are marked blue correspond to matched “y” ions. The number paired with each ion identification indicates the number of amino acids present. **(d)** HEK293T cells were transfected with Flag-tagged full-length wild-type Cdc14A, its inactive Cdc14A(PD) form or the indicated Cdc14A(PD) mutants. At 24 hours posttransfection, cells were treated with nocodazole during 12 hours and cellular extracts were obtained, resolved in Phostag-gels and analyzed by immunoblotting with anti-Flag antibodies. **(e)** Recombinant GST-Cdc14A(PD) and the different phosphorylation mutants (S453A, S411/549A, S411/549/453A (3A), S411/549/453/511A (4A) and S411/549/453/511/T590A (5A)) were phosphorylated *in vitro* by Cdk1-Cyclin B1 complexes in the presence of γ(^32^P)ATP. These results are representative from at least two different experiments.

Finally, we analyzed the ability of Cdk1-Cyclin B1 complexes to phosphorylate the different recombinant GST-Cdc14A(PD) mutants *in vitro*. We found that while Cdk1 efficiently phosphorylated Cdc14A(PD), the mutation of these three serines (S411/453/549A, 3A mutant) drastically reduced the level of Cdc14A(PD) phosphorylation (Fig. 4e). Mutation of the two additional Cdk1 consensus sites on the carboxyl-domain of Cdc14A (5A mutant) did not significantly further reduce the basal phosphorylation levels (Fig. 4e), indicating that *in vitro* serines 411, 453 and 549 are also targets of Cdk1. Amino acid sequence alignment among diverse Cdc14 phosphatase orthologues indicates the lack of conservation of these phosphorylation residues, specific for mammals (Supplementary Figure S3).

### Phosphorylation of Cdc14A by Cdk1 does not inhibit its catalytic activity

Given that human Cdc14A negatively regulates Cdk1 activity at the G2/M transition ^26,28^, coinciding with the Cdc14A hyperphosphorylation peak, we considered the possibility that mitotic phosphorylation might negatively regulate its catalytic activity, halting its inhibition of Cdk1 during early mitotic stages. In fact, it has been demonstrated that in the fission yeast *S. pombe* phosphorylation of Flp1 by Cdk1 during early mitosis reduces its catalytic activity to ensure proper progression through mitosis ^30^. Based on this finding, we evaluated the contribution of Cdk1 phosphorylation on the catalytic activity of human Cdc14A. Immunopurified Cdc14A from asynchronous or mitotic U-2-OS-Flag-Cdc14A cells was analyzed for its ability to hydrolyze *in vitro* the universal phosphatase substrate pNPP (4-Nitrophenyl phosphate disodium). We found that the rate at which Cdc14A samples hydrolyzed pNPP was similar in both asynchronous and mitotic samples (Supplementary Figure S4). Given that Cdc14A is able to autodephosphorylate, it could be possible that mitotic phosphorylations were autoremoved during the reaction and, in turn, the activity of Cdc14A was restored to interphase levels. To exclude this possibility, we examined the activity of Cdc14A phosphorylation mutants, Cdc14A-3A and Cdc14A-3E (a phosphomimetic version in which the three serines are replaced by glutamic acid), immunopurified from mitotic cells. We observed that Cdc14A-3A activity was similar to that of the wild-type, as would be expected if phosphorylation negatively regulated Cdc14A activity (Supplementary Figure S4). However, the rate at which the phosphomimetic mutant hydrolyzed pNPP was not reduced compared to the wild-type protein (Supplementary Figure S4). Although we cannot exclude the possibility that phosphomimetic mutant could fail to reproduce original phosphorylations ^40^, these data suggest that mitotic phosphorylation of Cdc14A does not affect its catalytic activity.

Next, we determined the activity of Cdc14A phosphorylation mutants *in vivo* using Wee1 kinase as substrate ^26^. HEK293T cells were co-transfected with HA-Wee1 and different forms of Flag-Cdc14A (WT, PD, 3A or 3E) and the effect on Wee1 was analyzed by changes in its electrophoretic mobility. As previously reported, Cdc14A, but not its catalytically inactive form, was able to dephosphorylate Wee1, as observed by its faster migration ^26^; (Fig. 5a). When cell cultures were enriched in mitosis by nocodazole treatment, expression of Cdc14A also affected the mobility of Wee1 and a band of lower molecular weight was generated (Fig. 5a), which also suggests that mitotic Cdc14A is not inhibited. Moreover, both phosphorylation mutant 3A and 3E showed a similar effect (Fig. 5a), also suggesting that mitotic phosphorylation does not reduce Cdc14A activity. To avoid the possibility that additional Cdk1 phosphorylation sites, although not identified under our experimental conditions, could be involved in the regulation of Cdc14A activity, we tested the activity of the nonphosphorylatable Cdc14A-5A and 7A mutants, where the five Cdk1 sites lying in the C-terminal domain or all seven ones were changed to Ala, respectively. These mutants did not show a greater effect on Wee1 when expressed in mitosis (Fig. 5a), and the corresponding phosphomimetic versions were not less active either (Fig. 5b). The same results were obtained when using KIBRA (kidney and brain expressed protein) as substrate ^21,41^ (Supplementary Figure S4), suggesting that Cdc14A activity was not compromised by phosphorylation on these residues.

**Figure 5.**
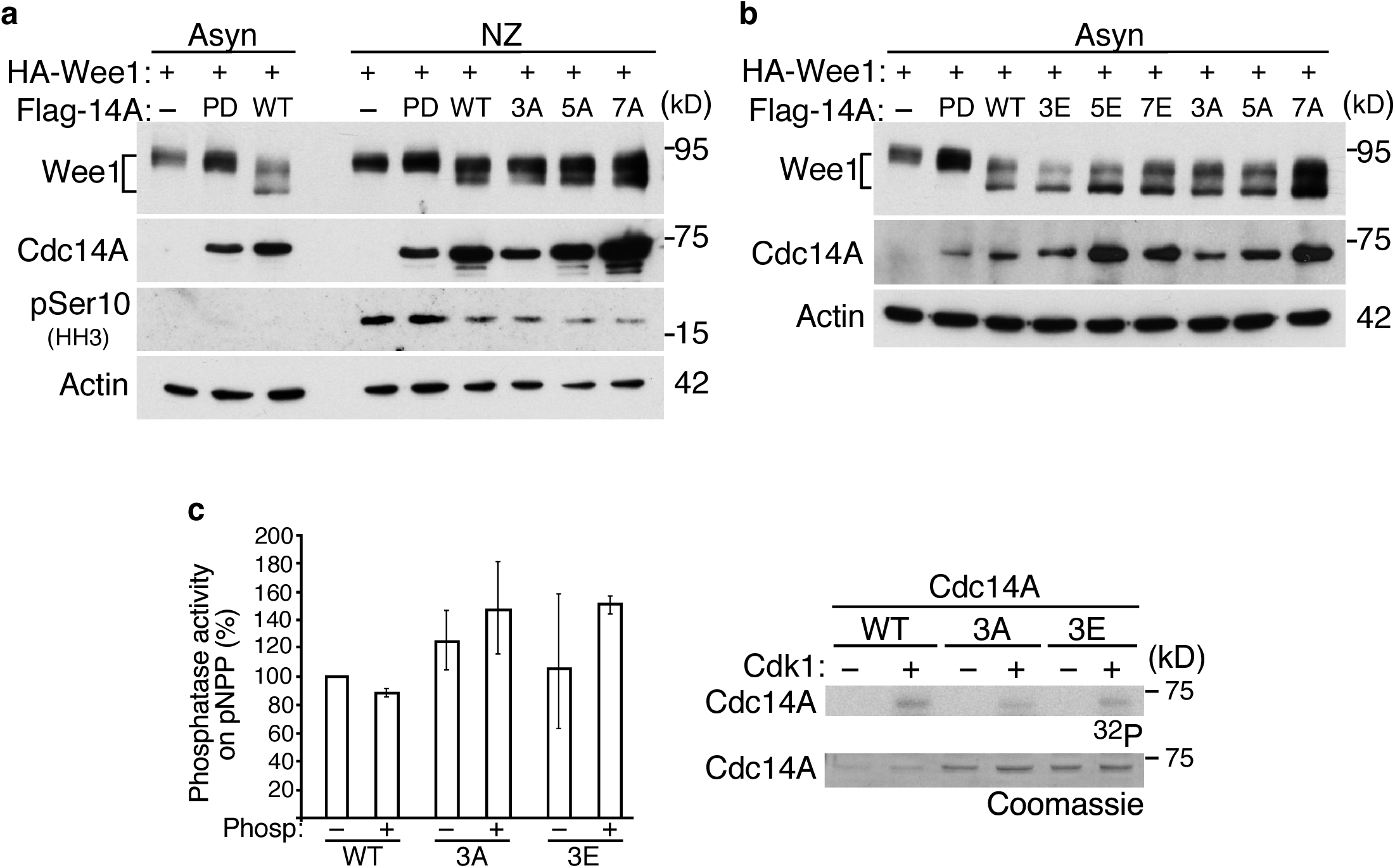
Cdc14A phosphorylation by Cdk1 does not inhibit its phosphatase activity. **(a,b)** HEK293T cells were co-transfected with HA-Wee1 and Flag-Cdc14A, the inactive Cdc14A(PD) form or the indicated Flag-tagged Cdc14A mutants. After 12 hours of transfection, half of the cells were treated with nocodazole during 12 hours. Then, cellular extracts were obtained from both asynchronously growing or nocodazole treated cells, and analyzed by immunoblot with the indicated antibodies. **(c)** GST-Cdc14A wild type (WT), GST-Cdc14A3A (3A) and GST-Cdc14A3E (3E) were purified from *E. coli* and processed with the PreScission protease to remove the tag. Samples were then incubated in the presence of Cdk1-Cyclin B1 complexes an cold-nonhydrolyzable ATP/γ(^32^P)ATP (100:1) and assayed for activity on pNPP. Reactions were performed in duplicate. Data correspond to the average of two different experiments. Protein levels were determined by coomassie and phosphorylation reactions by autoradiography (right panel is representative of one of these experiments).

To confirm that Cdk1 phosphorylation does not inhibit Cdc14A phosphatase activity, we performed *in vitro* kinase/phosphatase assays using purified recombinant Cdc14A proteins. To avoid the possibility that Cdc14A autocatalytically removes Cdk1 phosphorylation, we performed the kinase reactions using non-hydrolyzable ATP analog. After incubation with Cdk1, the ability of Cdc14A to hydrolyze pNPP was not significantly altered (Fig. 5c). The same was observed for Cdc14A-3A and Cdc14A-3E mutants: no difference between phosphorylated and non-phosphorylated protein activities (Fig. 5c). However, the activity of the two mutants appears slightly increased compared to the WT; since our previous results show that Cdk1 phosphorylation does not affect the activity of Cdc14A, small conformational changes due to the mutations could be responsible for it. Taken together and in concordance with a previous report ^29^, we conclude that Cdk1-mediated phosphorylation of human Cdc14A does not negatively regulate its catalytic activity.

### Cell cycle-regulated localization of Cdc14A is independent of Cdk1 phosphorylation

Human Cdc14A concentrates in centrosomes during interphase, and diffuses all over the cell during mitosis ^18,29^. Since the hyperphosphorylation state of Cdc14A peaks at the G2/M transition, coinciding with its relocalization, we investigated whether the release of Cdc14A from centrosomes depends on Cdk1 phosphorylation. If this were the case, we would expect the phosphomimetic mutant Cdc14A-3E to have less or no capacity to be retained in the centrosomes. Using U-2-OS cells transiently transfected to express EGFP-Cdc14A, EGFP-Cdc14A-3E or EGFP-Cdc14A-3A, we observed that during interphase EGFP-Cdc14A-3E localized to both the centrosome (determined by γ-tubulin staining) and the cytoplasm with the same pattern as wild type and the nonphosphorylatable mutant EGFP-Cdc14A-3A (Supplementary Figure S5). Furthermore, we examined EGFP-Cdc14A-3A and 3E staining during mitosis and found that their distribution was similar to that of wild type one; it appeared diffuse all over the cell, avoiding the chromatin, and was still present at the spindle poles, although to a lesser extent than in interphase centrosomes (Supplementary Figure S6). Thus, nor the lack of nor the constitutive phosphorylation resulted in a higher retention of Cdc14A in centrosomes during mitosis compared with the wild-type protein. These data suggest that Cdk1 phosphorylation is not necessary, at least by itself, to release Cdc14A from the centrosome at the onset of mitosis.

### Cdk1 phosphorylation regulates the binding pattern of Cdc14A at mitosis

Cdc14A protein levels show little or no variation throughout the cell cycle ^29^. We also found that the protein turnover of Cdc14A and its phosphorylation mutants (Cdc14A-3A and Cdc14A-3E) were quite similar (Supplementary Figure S7), suggesting that Cdk1 phosphorylation does not regulate the protein levels of Cdc14A during an unperturbed cell cycle. Thus, since the catalytic activity, subcellular localization and protein turnover of the Cdc14A phosphorylation mutants are similar to those of the wild-type protein, we can conclude that phosphorylation of these Cdc14A residues likely regulate specific and maybe still unknown functions.

Since the regulation of proteins by phosphorylation is often based on changes in its protein interaction pattern, we tested this possibility. We first checked how the phosphorylation state of Cdc14A affects the interaction with some of its known substrates or regulators, those other than Wee1 or KIBRA, with which the interaction does not seem to depend on phosphorylation (Fig. 5a,b and Supplementary Figure S4). It has been reported that Cdh1, the activator of APC/C (Anaphase-promoting complex/Cyclosome) during late mitosis, is a substrate of Cdc14A ^42^; therefore, we studied the Cdc14A-Cdh1 interaction by co-immunoprecipitation from cellular mitotic extracts. We detected that Cdh1 only interacts with the inactive form of Cdc14A, probably because interaction with the active form is transient and therefore difficult to detect. Our results show that both Cdc14A(PD) and the non-phosphorylatable Cdc14A(PD)-3A mutant bind equally to Cdh1 (Supplementary Figure S8). In addition, given that Plk1 has been shown to phosphorylate Cdc14A during mitosis ^34^, we explored whether or not Cdk1-mediated phosphorylation is needed to prime Plk1 phosphorylation on Cdc14A. Again, it was found that the interaction of Cdc14A with Plk1 during mitosis was not significantly reduced when the Cdc14A-3A mutant was expressed (Supplementary Figure S8), which agrees with our previous results showing that phosphorylation state does not have an impact on Cdc14A activity. In order to test whether or not Cdk1 phosphorylation could affect the binding of Cdc14A with other still unknown proteins, both Flag-tagged Cdc14A and Cdc14A-3A forms were transiently expressed in HEK293T cells and purified by immunoprecipitation from asynchronously growing or mitotic enriched cultures. Then, co-immunoprecipitates were resolved and identified on silver stained SDS-PAGE gels. As shown in Fig. 6a, some Cdc14A co-precipitated bands were detected in asynchronous samples with identical patterns in both the wild-type and Cdc14A-3A mutant. Moreover, additional proteins co-precipitated with wild-type Cdc14A when cultures were enriched in mitosis (Fig. 6a,b), and were less or not represented in Cdc14A-3A immunoprecipitates, indicating that Cdc14A interacts with a specific set of proteins at G2/M and/or early mitosis in a Cdk1-dependent manner. Next, in order to identify which proteins differentially interact with phosphorylated versus non-phosphorylated Cdc14A, the four bands identified as preferably bound to the phosphorylated form (Fig. 6a) were subjected to LC-MSMS analysis. In total, 121 proteins were identified by at least three unique peptides. Proteins likely to be nonspecific interactors (background contaminants), based on the CRAPome program, were identified and removed. A total of 33 Cdc14A-associated proteins were identified (Fig. 6c), of which 19 are phosphoproteins. Based on the GO annotation of their function, proteins are linked to biological processes related with metabolism (12), cell-cell adhesion and cell polarity (6), cell division (3), intracellular and intraflagellar transport (4), transcription (2) and others. None of them corresponds to known Cdc14A interactors.

**Figure 6.**
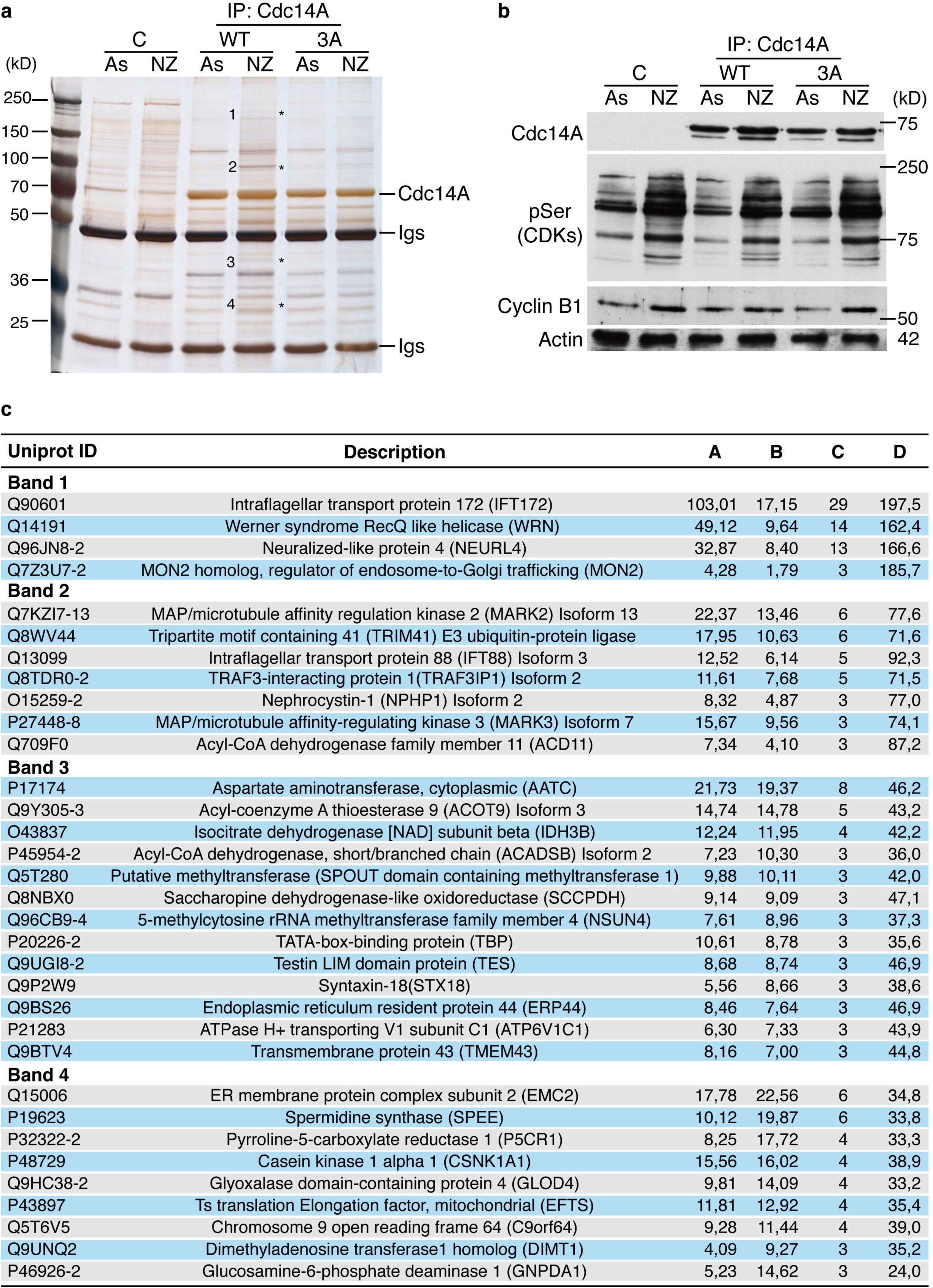
Cdk1-mediated Cdc14A phosphorylation modulates its binding pattern during mitosis. (a) Flag-Cdc14A or Flag-Cdc14A3A mutant expressed in HEK293T cells were immunoprecipitated with anti-Flag antibodies from asynchronously growing or nocodazole treated cells. Immunoprecipitates were resolved on SDS-PAGE gels and stained with silver solutions. As a negative control (C), extracts from HEK293T cells transfected in parallel with an empty Flag-vector were subjected to the same immunoprecipitation protocol and conditions. Asterisks show mitotic specific bands differently bound to the WT and the 3A mutant. **(b)** Cellular extracts used in (a) were analyzed by immunoblotting with the indicated antibodies. **(c)** Table showing Cdc14A-interacting proteins identified by LC-MSMS. Co-immunoprecipitates detected by silver staining in Fig. 6a (bands 1, 2, 3 and 4) were subjected to LC-MSMS analysis. From each of the four bands, all the proteins shown were identified with at least 3 unique peptides and selected from a total of 121 proteins after removing background contaminant proteins by using CRAPome program (contaminant repository for affinity purification; https://www.crapome.org).

Together, all these data suggest that phosphorylation of Cdc14A during early mitosis regulates its interaction with putative, but so far unknown, substrates or regulators. Future work is needed to confirm these Cdc14A interactions, which will help to elucidate the biological meaning of these phosphorylations and to discover new functions for human Cdc14A phosphatase.

## Discussion

Although the role of yeast Cdc14 in counteracting the activity of Cdk1 at the end of mitosis is well characterized, as are the molecular mechanisms for the regulation of its activity, both functions and regulatory mechanisms of human Cdc14 phosphatases are poorly understood. In this study, we characterized the human Cdc14A isoform as a substrate of Cdk1 at the entry into mitosis. Here, we report that Cdc14A is partially phosphorylated throughout S and G2 phases, becomes hyperphosphorylated at the entry into mitosis and is dephosphorylated as cells exit from mitosis, reaching G1 phase in a fully dephosphorylated state. Similar cell cycle phosphorylation kinetics has been observed for the fission yeast Flp1 orthologue^30^ and *Xenopus* Cdc14a^43^. In the former, phosphorylation by Cdk1 inhibits Flp1 catalytic activity until the exit from mitosis when autodephosphorylation restores its full phosphatase activity ^30^. In the case of the *Xenopus* Cdc14a isoform the meaning of its phosphorylation during early mitosis is not yet understood ^43^.

We have demonstrated by mass spectrometry and site-directed mutagenesis that Cdc14A is phosphorylated by Cdk1 at S411, S453 and S549, which are minimal Cdk consensus phosphorylation sites (Ser/Thr-Pro) laying on its carboxyl domain. The identification of these three major phosphoresidues allowed us to generate specific phosphorylation mutants for studying the biological meaning of this mitotic modification.

Since Cdc14A negatively regulates Cdk1 activity at the G2/M transition ^26,28^, we considered the possibility that Cdk1 phosphorylation at the early mitosis negatively regulates Cdc14A catalytic activity. Based on *in vitro* phosphatase assays, it has been reported that the enzymatic activity of Cdc14A does not significantly change during the cell cycle ^29^. Since Cdc14A has autodephosphorylation capacity, it could be possible that it reversed the potential phosphorylation-mediated inhibition of its activity during *in vitro* reactions. To discard this possibility, we performed *in vitro* phosphatase assays with the different Cdc14A phosphorylation mutants, and found that both the nonphosphorylatable and the phosphomimetic mutants had a catalytic activity similar to the wild-type protein. Similar results were obtained when the activity of Cdc14A mutants was analyzed *in vivo* using Wee1 kinase and KIBRA as endogenous substrates ^26,41^. Although we can not exclude that phosphomimetic mutations could fail to reproduce original phosphorylations, these results indicate that phosphorylation by Cdk1-Cyclin B1 complexes at early mitosis does not regulate human Cdc14A phosphatase activity.

Furthermore, this evidence shows that although Cdc14A shares the same Cdk1-mediated phosphorylation dynamics as its orthologue Flp1, the biological consequences of this modification are not the same. It has been reported that phosphorylation by Cdk1 also has a negative effect on the catalytic activity of Cdc14 in budding yeast. In this case, the S phase specific CDK complexes, Clb5-Cdc28, phosphorylate Cdc14 in its disordered tail to decrease its activity during S phase, which most likely modulates the balance of CDK and phosphatase activity ^11^. All these data support the idea that Cdk1 phosphorylation regulates Cdc14 in a different manner depending on the organism.

Another point of divergence between Flp1 and Cdc14A orthologues is that although both have autophosphatase activity, Cdc14A does not fully autoremove Cdk1 phosphorylation at the end of mitosis like Flp1 does ^30^. The PPP family members PP2A and PP1 have prominent roles in dephosphorylating Cdk1 substrates at this cell cycle stage in mammals ^37,38,44^. We have shown that Cdc14A interacts *in vivo* with PP2A-Cα, suggesting that PP2A may be the phosphatase that dephosphorylates it. Further work will be required to explore this possibility.

As in yeast, human Cdc14 localization changes during the cell cycle. Thus, Cdc14A, concentrated in centrosome during interphase, is distributed all over the cell in mitosis ^16^. Different regions of Cdc14A, including the C-terminal domain, are involved in its centrosome localization ^18^. Our data suggest that Cdc14A centrosome release at the G2/M transition is independent of phosphorylation by Cdk1 at its carboxyl terminus. Whether Cdk1 phosphorylation could act in concert with other phosphorylations or additional modifications to regulate Cdc14A localization and/or catalytic activity, is not yet known.

Finally, we considered the possibility that Cdk1-mediated Cdc14A phosphorylation might play such a regulatory role, contributing to determine the specific Cdc14A protein interaction landscape during this cell cycle window. Consistent with this idea, immunoprecipitation analysis showed that some proteins associate with Cdc14A specifically during mitosis, and that these interactions are Cdk1 phosphorylation dependent. The interactions studied, focused on some of its already characterized substrates or regulators (Cdc14A-Wee1, -Cdh1, -KIBRA and -Plk1) proved, however, to be independent of the Cdc14A phosphorylation state, suggesting that other proteins, likely not known yet, are involved in these interactions. We have identified a pool of potential Cdc14A-associated proteins that bounds Cdc14A in a Cdk1-dependent phosphorylation manner. Although none of them corresponds to known Cdc14A interactors, some are linked to biological process in which Cdc14A has been already involved, such as cell polarity ^21^, cell division ^18,26,28^, and transcription ^23,25^. Future work is needed to confirm these interactions, which will contribute to further understand both the regulatory mechanisms of the human Cdc14A isoform and its still poorly characterized functions, especially those related with mitosis. Different research studies have pointed out that yeast Cdc14 functions are not fully conserved in other organisms (reviewed by ^16^). Our work provides new data about the regulation of human Cdc14A by Cdk1 phosphorylation, which also differs from yeast.

## Methods

### Cell culture, synchronization and drug treatment

Derivatives of the U-2-OS and HeLa cell lines expressing HA-Cdc14A or Flag-Cdc14A alleles, respectively, were generated by infection with specific retroviruses produced in HEK293T. RPE-Cdc14A^-/-^ cells were a kind gift from the laboratory of Dr. P. Jallepalli ^16^. U-2-OS, HeLa and HEK293T cells were cultured in Dulbecco Modified Eagle’s Medium (DMEM) supplemented with 2mM glutamine, 100U/ml penicillin G, 0.1mg/ml streptomycin and 10% fetal bovine serum at 37 °C/5% CO_2_. HEK293T and U-2-OS cells were transfected with the calcium phosphate/HEBS (HEPES-buffered saline solution) method or Lipofectamine 2000 (GIBCO Invitrogen), respectively. U-2-OS cells were synchronized in G1/S phase by double thymidine treatment. Briefly, thymidine, 2.5mM, was added for 24 hours, followed by a release period of 10 hours and a second treatment with thymidine for a further 24 hours. Then, the cells were released to fresh culture medium after extensive washing with PBS. Where indicated, nocodazole (50ng/ml) was added to avoid exit from mitosis. Mitotic cells were obtained by shaking off the rounded cells following 12 hours of treatment with nocodazole (50ng/ml). HEK293T cells were enriched in mitosis by a 12 hr treatment with nocodazole. When indicated, cells were treated with 10µM purvalanol A and 10µM RO-3306 (Calbiochem), 0.5µM Okadaic acid (Sigma). For estimation of the protein half-life, the cultures were treated with 25µg/ml Cycloheximide (Sigma) for the times indicated.

### Plasmids, mutagenesis and recombinant protein expression

Human Cdc14A cDNA was cloned into pCEFL-HA, pCEFL-Flag, pCDNA3-Myc or pEGFP-N1 mammalian expression vectors using suitable restriction enzymes. Two truncated constructions corresponding to the N-amino domain (aas 1–332) and C-carboxyl domain (aas 333–623) were generated by polymerase chain reaction (PCR) using the corresponding primers. Cdc14A phosphorylation mutants were generated using site-directed mutagenesis. For viral infection, wild-type and mutant Cdc14A cDNAs were subcloned into pBabe-puro retroviral expression vector. Retroviruses were produced in HEK293T cells and used to infect U-2-OS or HeLa cells. Infected cells were selected by adding puromycin (0.5 µg/ml) to the culture medium. pBabe-puro-EGFP plasmid was used to control infection efficiency. For bacterial expression in *E. coli* and the subsequent purification of recombinant GST-proteins, wild-type and mutant Cdc14A/B constructs were subcloned into a pGEX-4T-GST or pGEX-6T-GST vectors. GST-recombinant proteins were purified on glutathione agarose beads using as lysis buffer PBS containing 1% Triton X-100 and supplemented with complete protease inhibitor mixture (Roche). Where indicated, treatment with PreScission protease (Ref. 270843, Amersham GE helthcare) was used following the manufacturer’s instructions to remove GST-tag. All the material generated during the current study is available from the corresponding author upon request.

### Flow cytometry

For cell cycle analysis, cells were fixed in ice-cold 70% PBS-ethanol and stained with PBS containing 8 µg/ml propidium iodide and 20 µg/ml-RNaseA for 1 hour at 37 °C. For phospho-histone H3 positivity, cells were harvested and fixed in ice-cold 70% ethanol. Cells were washed with PBS-0.15% Triton X-100 and subsequently stained with rabbit phospho-histone-H3(S10) (Ref. 06–570, Millipore; 1:200) and Fluorescein isothiocyanate-conjugated goat anti rabbit IgG (Ref. 111-095-144, lot 79638, Jackson Immunoresearch Lab. INC; 1:200) antibodies. Then, they were incubated with 8 µg/ml-propidium iodide and 20 µg/ml-RNase A solution for 1 hour at 37 °C. Stained cells were acquired using a FACSCalibur device (Becton Dickinson) and cell cycle distribution was analysed using Cell Quest Pro.

### Immunochemical techniques

For western blotting, cells were harvested and lysed in buffer containing 0.5% NP40, 20mM Tris-Cl pH8.0, 150mM NaCL, 1mM DTT, 10mM β-glycerophosphate, 5mM NaF, 1mM Na_3_VO_4_ and supplemented with complete protease inhibitor mixture (Roche). Protein samples were run on 8, 10 or 12% SDS-acrylamide gels. For phosphorylation analyses, Phostag (Wako Lab. Chem.) and MnCl_2_ (100mM) were added to the SDS-acrylamide mix following the manufacturer’s instructions. The primary antibodies used were mouse HA-tag (12CA5, Ref. 11-666606001, Boehringer Mannheim), mouse Flag-tag (M2, peroxidase conjugated, Ref. A8592, Sigma) or Rat anti-Flag, (Ref. 299474, Agilent Technologies), mouse Myc-tag (9E10, Ref. M5546, Sigma), mouse Cdk1 (Ref. sc-54, Santa Cruz Biotechnology), rabbit phospho-Tyr15-Cdk1 (Ref. 9111, Cell Signalling), rabbit phospho-(Ser) CDKs (Ref. 2324, Cell Signaling), goat Cdc14A (N-18, Ref. sc-25952, Santa Cruz Biotechnology), rabbit Cyclin B1 (Ref. sc-752, Santa Cruz Biotechnology), mouse PP2A (ID6, Ref. 05-421, Millipore), rabbit Wee1 (Ref. sc-9037, Santa Cruz Biotechnology), mouse Plk1 (Ref. 33-1700, Zymed), rabbit phospho-histone-H3(S10) (Ref. 06-570, Millipore) and mouse β-Actin (AC-15, Ref. A5441, Sigma). Specificity of anti-HA and anti-Flag antibodies were checked using control wild type U-2-OS or HEK293T cells that do not express the corresponding tagged-Cdc14A forms. For the immunoprecipitation studies, cell lysates (0.5, 1 or 5mg of protein) were incubated with the corresponding antibodies (2µg/mg protein extracts) and protein G Dynabeads (Ref 100-04D, Invitrogen) (10µl/mg protein) for 3–4 hours at 4 °C. Immunoprecipitates were collected by centrifugation, washed five times with lysis buffer, eluted by boiling with Laemli buffer at 95 °C for 5 min and subjected to SDS-PAGE electrophoresis and immunoblot analysis.

### Kinase and phosphatase assays

All recombinant GST-proteins produced in bacteria were purified on glutathione beads (GST). Approximately 0.5mg of recombinant GST-Cdc14A, GST-Cdc14B, GST-Cdc14A(PD) or GST-Cdc14B(PD) were phosphorylated by Cdk1-Cyclin B1 (New England Biolabs) at 30 °C in the presence of 0.10 µCi of γ(^32^P)ATP with or without sodium vanadate (1mM) for 30 minutes.

For phosphatase assays, recombinant GST-Cdc14A(PD) was phosphorylated by Cdk1-Cyclin B1 in the presence of 0.10 µCi of γ(^32^P)ATP for 30 minutes at 30 °C. Beads were washed extensively in phosphatase buffer (20 mM Tris, pH 8.3; 150 mM NaCl; 2 mM EDTA; 0.1% Triton X-100; 5 mM DTT) to remove Cdk1 and non-incorporated γ(^32^P)ATP, and then incubated in the mentioned phosphatase buffer in the presence of recombinant GST-Cdc14A, GST-Cdc14B, GST-Cdc14A(PD) or GST-Cdc14B(PD) at 30 °C for 30–45 min. Reactions were stopped by the addition of sample buffer and boiled for 5 minutes at 95 °C. Samples were then resolved on SDS-PAGE and visualized by Coomassie staining and autoradiography. Where indicated, recombinant GST-Cdc14A forms were purified from *E. coli* and processed with the PreScission protease (Amersham GE healthcare) following the manufacturer’s instructions to remove the tag. Samples were then incubated in the presence of Cdk1-Cyclin B1 complexes an cold-nonhydrolyzable ATP/γ(^32^P)ATP (100:1) and assayed for activity on pNPP.

For phosphatase assays on pNPP (4-Nitrophenyl phosphate disodium; Sigma), Flag-Cdc14A immunoprecipitates were resuspended in phosphatase buffer to 100mg/200ml in 96-well plates. pNPP was added to a final concentration of 5mM and absorbance at 405 nm was monitored at 30 °C for 30 minutes every 5 minutes. Reactions were made in duplicate or triplicate as stated. Absorbance readings were plotted and the rates of the reactions were determined using linear regression analysis, where the slope was used as indicative of the reaction speed and therefore the activity of the protein. The 25% of the immunoprecipitates was detected by Western blot and quantified for normalization.

### Immunofluorescence Microscopy

U-2-OS-HA-Cdc14A cell lines were seeded onto sterile, Poly-L-Lysine (Sigma) -treated coverslips in 6-well plates, fixed with paraformaldehyde (4%) and permeabilized with methanol. Then, cells were blocked in PBS with 1% bovine serum albumin (BSA) (Sigma) and incubated in the same buffer with rabbit anti-HA (C29F4, Ref 3724, Cell signaling; 1:1000) and mouse anti-γ-tubulin (GTU-88, Ref. T6557, Sigma; 1:7500) antibodies for 1 hour. Cells were washed three times with PBS/1% BSA and incubated with Fluorescein isothiocyanate-conjugated goat anti rabbit IgG (Ref. 111-095-144, lot 79638, Jackson Immunoresearch Lab. INC) and Cy™3-conjugated goat anti-mouse IgG (Ref. 115-165-003, lot 112660, Jackson Immunoresearch Lab. INC), respectively, for 30 minutes to visualize the primary antibodies. Staining specificity of anti-HA antibodies was checked using control wild type U-2-OS cells. No staining signal was detected in these cells. To detect EGFP-Cdc14A, U-2-OS cells were transfected using lipofectamine 2000 reagents (Invitrogen) with the different pEGFP-N1-Cdc14A DNA plasmids. After 36 hr of transfection, cells were fixed and permeabilized as above indicated, and incubated with mouse anti-γ-tubulin antibody (GTU-88, Ref. T6557, Sigma; 1:7500). Cells were washed three times with PBS and DNA was stained with DAPI. Images were acquired using a Leica DM6000B or a Zeiss Axioimager Apotome microscopes (objective 40X) and processed using OpenLab 4.0.3 or Zeiss Zen lite softwares.

### Mass Spectrometry LC-MS/MS analysis and database search

Immunoprecipitates of Myc-Cdc14A and Myc-Cdc14A(PD) from untreated or Okadaic acid-treated HEK293T cells (0.5µM OA during 2 hours) were resolved on SDS-PAGE gels and silver stained. Gel bands were processed as reported ^45^ using a 1:50 trypsin:protein ratio; the peptide mixture was analyzed by LC-MS/MS in an nanoAcquity UPLC (Waters Corp., Milford, MA) coupled with an LTQ - Orbitrap Velos (Thermo-Fisher, San Jose, CA). Peptides were trapped and separated by a 30 min gradient to 35% B in a Symmetry and BEH C18 columns (Waters Corp., Milford, MA). A TOP 20 followed by CID and TOP 10 followed by CID MSA data-dependent methods were used for peptide and CDC14 phosphopeptide analysis respectively. Protein identification was done using Sequest algorithm and *Homo sapiens* RefSeq database release January 2014 from NCBI including common contaminant sequences. Percolator was used for peptide validation. In addition, phosphopeptides were confirmed by manual interpretation of MSA ion spectra.

## Acknowledgments

We thank S. Andrés for technical assistance and other members of laboratory for helpful discussions. We are grateful to Dr. I. García-Higuera and S. Moreno (IBFG, Salamanca) for Cdh1 plasmids and anti-Plk1 antibodies, Dr. J. Dong (Univ. Nebraska Medical Center, Nebraska) for the KIBRA plasmids, Dr. C. Guerrero (IBMCC, Salamanca) for the anti-PP2A antibody and Dr. Jallepalli (Memorial Sloan Kettering Cancer Center, New York) for the RPE Cdc14A^-/-^ cells. We are grateful to the proteomics facility of Centro de Investigación del Cáncer, Salamanca, Spain, where the proteomic analysis was performed, Grant PRB2 (IPT13/0001 - ISCIII-SGEFI / FEDER). This work was funded by grants from the Spanish Ministry of Economy, Industry and Competitiveness (MINECO; BFU2015-69709-P and SAF2015-69920-R). S.O. was supported by a FPU fellowship from the Spanish Ministry of Education and P.A. was supported by a JAE-Predoctoral fellowship from the Spanish National research Council (CSIC).

## Author contributions

S.O., A.B. and M.P.S. conceived the study and analysed the data with critical inputs from M.M. and F.X.P-M. P.A. generated Cdc14A mutants and S.O. performed most of the experiments. M.P.S wrote the paper and all authors reviewed and discussed the results and approved the final version of the manuscript.

## Additional Information

### Supplementary information

#### Competing Interests

The authors declare that they have no competing interests.

## References

1 Bremmer, S. C. et al. Cdc14 phosphatases preferentially dephosphorylate a subset of cyclin-dependent kinase (Cdk) sites containing phosphoserine. J Biol Chem 287, 1662–1669 (2012).

2 Kuilman, T. et al. Identification of Cdk targets that control cytokinesis. EMBO J 34, 81–96, doi:embj.201488958 [pii] 10.15252/embj.201488958 (2015).

3 Meitinger, F., Palani, S. & Pereira, G. The power of MEN in cytokinesis. Cell Cycle 11, 219–228, doi:18857 [pii]. 10.4161/cc.11.2.18857 (2012).

4 Queralt, E. & Uhlmann, F. Cdk-counteracting phosphatases unlock mitotic exit. Curr Opin Cell Biol 20, 661–668 (2008).

5 Stegmeier, F. & Amon, A. Closing Mitosis: The Functions of the Cdc14 Phosphatase and Its Regulation. Annu Rev Genet (2004).

6 Clifford, D. M. et al. The Clp1/Cdc14 phosphatase contributes to the robustness of cytokinesis by association with anillin-related Mid1. J Cell Biol 181, 79–88 (2008).

7 Wolfe, B. A., & Gould, K. L. Fission yeast Clp1p phosphatase affects G(2)/M transition and mitotic exit through Cdc25p inactivation. Embo J 23, 919–929 (2004).

8 Esteban, V. et al. A role for the Cdc14-family phosphatase Flp1p at the end of the cell cycle in controlling the rapid degradation of the mitotic inducer Cdc25p in fission yeast. J Cell Sci 117, 2461–2468 (2004).

9 Cueille, N. et al. Flp1, a fission yeast orthologue of the s. cerevisiae CDC14 gene, is not required for cyclin degradation or rum1p stabilisation at the end of mitosis. J Cell Sci 114, 2649–2664 (2001).

10 Trautmann, S. et al. Fission yeast Clp1p phosphatase regulates G2/M transition and coordination of cytokinesis with cell cycle progression. Curr Biol 11, 931–940 (2001).

11 Eissler, C. L. et al. The Cdk/cDc14 module controls activation of the Yen1 holliday junction resolvase to promote genome stability. Mol Cell 54, 80–93, doi:S1097-2765(14)00128-2 [pii]. 10.1016/j.molcel.2014.02.012 (2014).

12 Villoria, M. T. et al. Stabilization of the metaphase spindle by Cdc14 is required for recombinational DNA repair. Embo J 36, 79–101 (2016).

13 Díaz-Cuervo, H. & Bueno, A. Cds1 controls the release of Cdc14-like phosphatase Flp1 from the nucleolus to drive full activation of the checkpoint response to replication stress in fission yeast. Mol Biol Cell 19, 2488–2499 (2008).

14 Vázquez-Novelle, M. D., Esteban, V., Bueno, A. & Sacristán, M. P. Functional homology among human and fission yeast Cdc14 phosphatases. J Biol Chem 280, 29144–29150 (2005).

15 Li, L., Ernsting, B. R., Wishart, M. J., Lohse, D. L. & Dixon, J. E. A family of putative tumor suppressors is structurally and functionally conserved in humans and yeast. J Biol Chem 272, 29403–29406 (1997).

16 Mocciaro, A. et al. Vertebrate cells genetically deficient for Cdc14A or Cdc14B retain DNA damage checkpoint proficiency but are impaired in DNA repair. J Cell Biol 189, 631–639 (2010).

17 Sacristán, M. P., Ovejero, S. & Bueno, A. Human Cdc14A becomes a cell cycle gene in controlling Cdk1 activity at the G(2)/M transition. Cell Cycle 10, 387–391, doi:14643 [pii] 10.4161/cc.10.3.14643 (2011).

18 Mailand, N. et al. Deregulated human Cdc14A phosphatase disrupts centrosome separation and chromosome segregation. Nat Cell Biol 4, 317–322 (2002).

19 Krasinska, L. et al. Regulation of multiple cell cycle events by Cdc14 homologues in vertebrates. Exp Cell Res 313, 1225–1239 (2007).

20 Wu, J. et al. Cdc14B depletion leads to centriole amplification, and its overexpression prevents unscheduled centriole duplication. J Cell Biol 181, 475–483 (2008).

21 Chen, N. P., Uddin, B., Voit, R. & Schiebel, E. Human phosphatase CDC14A is recruited to the cell leading edge to regulate cell migration and adhesion. Proc Natl Acad Sci U S A 113, 990–995, doi:1515605113 [pii] 10.1073/pnas.1515605113.

22 Lin, H. et al. Cdc14A and Cdc14B Redundantly Regulate DNA Double-Strand Break Repair. Mol Cell Biol 35, 3657–3668 (2015).

23 Guillamot, M. et al. Cdc14b regulates mammalian RNA polymerase II and represses cell cycle transcription. Sci Rep 1, 189 (2011).

24 Bassermann, F. et al. The Cdc14B-Cdh1-Plk1 axis controls the G2 DNA-damage-response checkpoint. Cell 134, 256–267 (2008).

25 Clemente-Blanco, A. et al. Cdc14 phosphatase promotes segregation of telomeres through repression of RNA polymerase II transcription. Nat Cell Biol 13, 1450–1456 (2011).

26 Ovejero, S., Ayala, P., Bueno, A. & Sacristan, M. P. Human Cdc14A regulates Wee1 stability by counteracting CDK-mediated phosphorylation. Mol Biol Cell 23, 4515–4525 (2012).

27 Tumurbaatar, I., Cizmecioglu, O., Hoffmann, I., Grummt, I. & Voit, R. Human Cdc14B promotes progression through mitosis by dephosphorylating Cdc25 and regulating Cdk1/cyclin B activity. PLoS One 6, e14711 (2011).

28 Vázquez-Novelle, M. D., Mailand, N., Ovejero, S., Bueno, A. & Sacristán, M. P. Human Cdc14A phosphatase modulates the G2/M transition through Cdc25A and Cdc25B. J Biol Chem (2010).

29 Kaiser, B. K., Zimmerman, Z. A., Charbonneau, H. & Jackson, P. K. Disruption of centrosome structure, chromosome segregation, and cytokinesis by misexpression of human Cdc14A phosphatase. Mol Biol Cell 13, 2289–2300 (2002).

30 Wolfe, B. A., McDonald, W. H., Yates, J. R., 3rd & Gould, K. L. Phospho-regulation of the Cdc14/Clp1 phosphatase delays late mitotic events in S. pombe. Dev Cell 11, 423–430 (2006).

31 De Wulf, P., Montani, F. & Visintin, R. Protein phosphatases take the mitotic stage. Curr Opin Cell Biol 21, 806–815 (2009).

32 Amon, A. A decade of Cdc14--a personal perspective. Delivered on 9 July 2007 at the 32nd FEBS Congress in Vienna, Austria. FEBS J 275, 5774–5784, doi:EJB6693 [pii]. 10.1111/j.1742-4658.2008.06693.x (2008).

33 Trautmann, S., Rajagopalan, S. & McCollum, D. The S. pombe Cdc14-like phosphatase Clp1p regulates chromosome biorientation and interacts with Aurora kinase. Dev Cell 7, 755–762 (2004).

34 Yuan, K. et al. Phospho-regulation of HsCdc14A By Polo-like kinase 1 is essential for mitotic progression. J Biol Chem 282, 27414–27423 (2007).

35 St-Denis, N. et al. Phenotypic and Interaction Profiling of the Human Phosphatases Identifies Diverse Mitotic Regulators. Cell Rep 17, 2488–2501 (2016).

36 Mochida, S., Ikeo, S., Gannon, J. & Hunt, T. Regulated activity of PP2A-B55 delta is crucial for controlling entry into and exit from mitosis in Xenopus egg extracts. EMBO J 28, 2777–2785, doi:emboj2009238 [pii] 10.1038/emboj.2009.238 (2009).

37 Schmitz, M. H. et al. Live-cell imaging RNAi screen identifies PP2A-B55alpha and importin-beta1 as key mitotic exit regulators in human cells. Nat Cell Biol 12, 886–893 (2010).

38 Manchado, E. et al. Targeting mitotic exit leads to tumor regression in vivo: Modulation by Cdk1, Mastl, and the PP2A/B55alpha, delta phosphatase. Cancer Cell 18, 641–654 (2010).

39 Thingholm, T. E., Jensen, O. N., Robinson, P. J. & Larsen, M. R. SIMAC (sequential elution from IMAC), a phosphoproteomics strategy for the rapid separation of monophosphorylated from multiply phosphorylated peptides. Mol Cell Proteomics 7, 661–671, doi:M700362-MCP200 [pii]. 10.1074/mcp.M700362-MCP200 (2008).

40 Dephoure, N., Gould, K. L., Gygi, S. P. & Kellogg, D. R. Mapping and analysis of phosphorylation sites: a quick guide for cell biologists. Mol Biol Cell 24, 535–542, doi:24/5/535 [pii]. 10.1091/mbc.E12-09-0677 (2013).

41 Ji, M. et al. Phospho-regulation of KIBRA by CDK1 and CDC14 phosphatase controls cell-cycle progression. Biochem J 447, 93–102 (2012).

42 Bembenek, J. & Yu, H. Regulation of the anaphase-promoting complex by the dual specificity phosphatase human Cdc14a. J Biol Chem 276, 48237–48242 (2001).

43 Kaiser, B. K., Nachury, M. V., Gardner, B. E. & Jackson, P. K. Xenopus Cdc14 alpha/beta are localized to the nucleolus and centrosome and are required for embryonic cell division. BMC Cell Biol 5, 27 (2004).

44 Qian, J. et al. Cdk1 orders mitotic events through coordination of a chromosome-associated phosphatase switch. Nat Commun 6, 10215 (2015).

45 Shevchenko, A., Wilm, M., Vorm, O. & Mann, M. Mass spectrometric sequencing of proteins silver-stained polyacrylamide gels. Anal Chem 68, 850–858 (1996).

